# Advancing ecological networks: moving beyond binary classification to probabilistic interactions

**DOI:** 10.1101/2024.02.07.579289

**Authors:** Martin Philippe-Lesaffre

**Affiliations:** Université Paris-Saclay, CNRS, AgroParisTech, Ecologie Systématique Evolution, IDEEV, 12 rue 128, Gif-sur-Yvette, France, 91190

## Abstract

Leveraging trophic interactions to deduce macro-ecological patterns has become a prevalent method, taking advantage of the extensive databases on binary trophic interactions (i.e., prey-predator relationships). However, this binary approach oversimplifies complex ecological dynamics and fails to capture the nuanced structure of food webs. The challenge lies in the scarcity and limited availability of data on non-binary interactions, which are crucial for a more comprehensive understanding of ecological networks. This study explores the use of binary classifiers, particularly the XGBOOST algorithm to address the limitations of traditional binary approaches to prey-predator relationships. By predicting predation probabilities among nine mammalian predators using species traits, my findings demonstrate the classifiers’ robust predictive capabilities to binary predictions but also a good correlation between probabilistic predation derived from binary classifiers and observed prey preferences. It also highlighted the importance of selecting informative species traits for predicting interaction, with performance contrastingly superior to null models. Despite a small sample size, this work provides insightful results and sets a foundation for future research to expand these models to broader ecological networks, emphasizing the need for comprehensive prey preference data.

## Introduction

Empirical records form the backbone of understanding trophic interactions, yet supervised learning combined with open databases increasingly aids in extrapolating unknown relationships, leveraging species traits and phylogeny (Llewelyn et al., 2022, 2023). However, a critical limitation in these studies lies in the binary coding of trophic interactions (0 for absence, 1 for presence), which can obscure the nuances and biomass fluxes intrinsic to each interaction. This simplification often results in overly simplistic food web topologies and consequently, less precise models (Gaüzère et al., 2022, 2023). The growing availability of databases cataloging species traits across various taxa, particularly for tetrapods, presents a rich resource (Oliveira et al., 2017; Soria et al., 2021; Tobias et al., 2022). Given that supervised learning methods, incorporating traits and phylogeny, have shown promise in predicting binary interactions (Caron et al., 2022; Desjardins-Proulx et al., 2017; Llewelyn et al., 2023), it is imperative to explore the potential of these methods for delineating more complex interactions.

These supervised learning approaches, using binary classifiers (e.g., generalized linear model, random forest, extreme gradient boosting, deep neural networks), not only predict binary outcomes but also offer probabilities, indicating the likelihood of a species being prey. This raises a crucial question: does this probability reflect the intensity of predation or the mere likelihood of a species being prey? To address this, I utilized mammalian prey preference data from Carnivora predators in Europe sourced from the CARNIDIET database (Middleton et al., 2021), and integrated it with trait data from combined databases (Soria et al., 2021). I then computed a predicted predation probability for each prey-predator pair by computing binary classification with optimized extreme gradient boosting (XGBOOST) models where prey preference was transformed into binary response, predation, or no predation. I selected XGBOOST as the primary modeling tool due to several compelling reasons. Foremost among these is its consistent performance; XGBOOST generally outperforms most tree-based and regression-based models in predictive accuracy. This superiority makes it an ideal choice for complex ecological data analyses where precision is paramount. Additionally, XGBOOST’s ease of implementation in R, as facilitated by the dedicated XGBOOST package (Chen & Guestrin, 2016), significantly streamlines the modeling process. This accessibility is crucial for ensuring that the model can be efficiently applied and replicated in ecological research. Moreover, the recent advancements in interpretability techniques, such as SHAP (SHapley Additive exPlanations; Lundberg & Lee, 2017), further enhance the appeal of XGBOOST. SHAP provides a robust framework for deciphering and communicating the contributions of individual predictors in the model, a feature that is invaluable in understanding and validating the ecological insights derived from the model. The combination of high performance, user-friendly implementation, and advanced interpretability techniques makes XGBOOST a superior choice for this study.

Assessing the predictive power of the classifiers, using the Matthews correlation coefficient, I compared its results with the observed prey preferences by calculating the Spearman’s rank correlation coefficient. To validate that traits are effective predictors of trophic interactions, I also performed null models for each predator.

## Methods

### Software and Version

All analyses in this study were conducted using R Studio, Version 2023.03.1+446 (R Core Team, 2022).

### Compiling Datasets of Prey-Predator Relationships

The diet data was extracted from the CARNIDIET database (Middleton et al., 2021) on December 20th, accessible at https://owen-middleton.shinyapps.io/CarniDIET-Shiny/. To streamline analysis, I focused on six aspects of the database: (i) predator species, (ii) prey species, (iii) country of study, (iv) GPS coordinates, (v) whether the study occurred on an island, and (vi) percentage of each species in the predator’s diet. Only mammal species were assessed at species level in CARNIDIET so I focused on this subset of the diet of Carnivora. The initial filtering included retaining only species-level identified prey, studies from Europe, and excluding island observations. This process resulted in predator-specific datasets, each containing species-level prey data, dietary percentages, and study locations. Datasets with fewer than 100 observations were excluded to mitigate sampling bias.

Further, all native mammals in Europe were collated from the IUCN Red List database (IUCN, 2023), focusing on those with ‘Native’ origin status and ‘Extant’ presence. Each predator dataset was then merged with this information, adding non-present mammal species with a dietary percentage set to zero. The geographical ranges of European mammals were sourced from the IUCN RedList, enabling the exclusion of species from the predator dataset whose ranges did not overlap with any study locations. For species appearing multiple times, I averaged the values to derive a singular data point per species. Seven traits were extracted from the combined database (Soria et al., 2021) and IUCN RedList: body mass, brain mass, foraging niche, habitat breadth, trophic level, activity period, and litter size.

This resulted in 10 datasets, corresponding to the predation preferences of 10 predators. A column was added to each dataset to categorize prey: ‘1’ if predation preference differed from 0, ‘0’ otherwise. Ursus arctos was finally removed because only 3 prey were found with prey preference. This lead to a dataset for *Canis aureus, Martes martes, Martes foina, Canis lupus, Vulpes vulpes, Felis sylvestris, Genetta genetta, Neovison vison* and *Lynx lynx* (Fig. 1).

**Figure 1.**
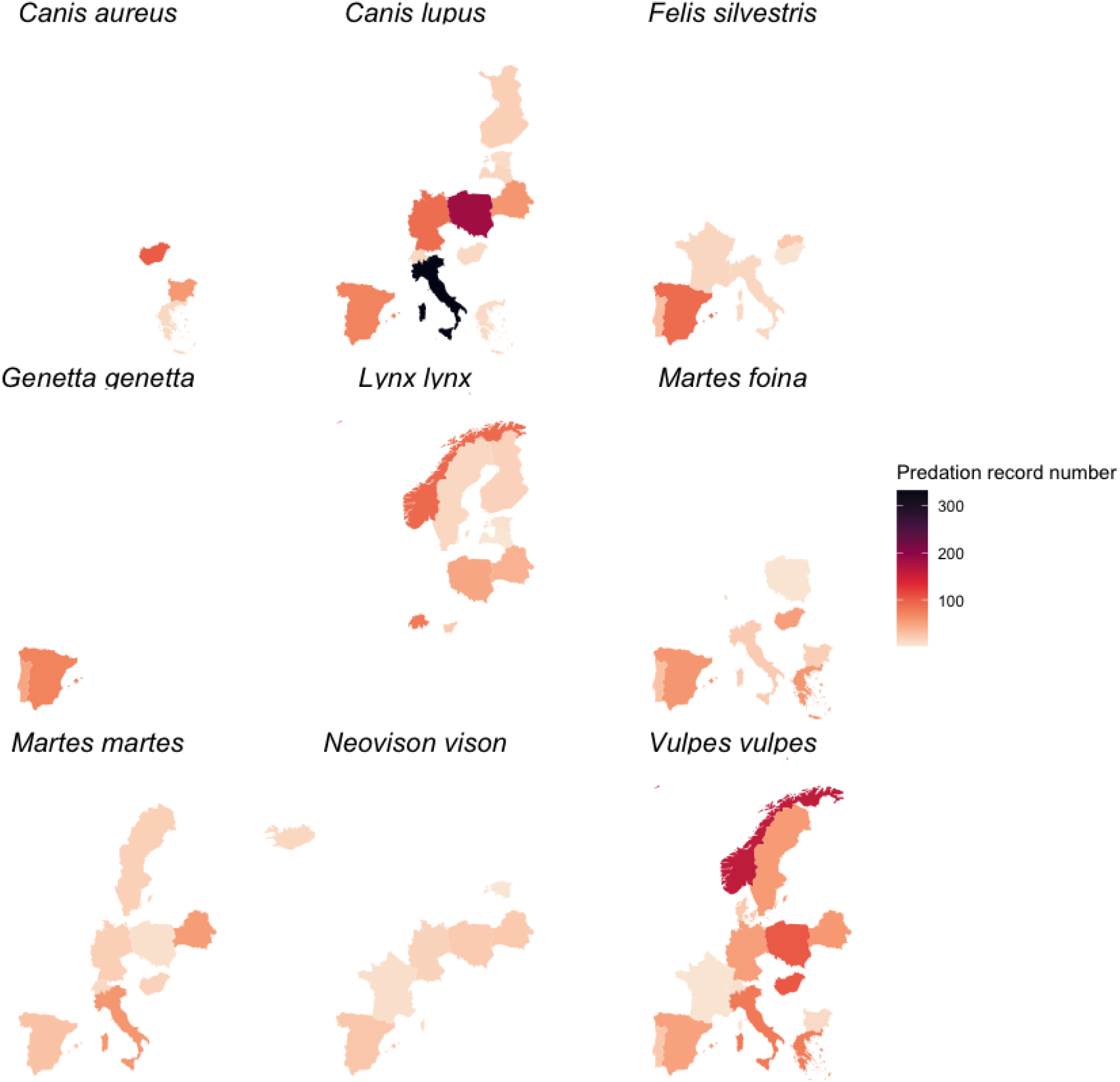
showcases the geographical distribution of mammalian prey predation records per predator, as identified at the species level and sourced from the CARNIDIET database (Middleton et al., 2021). Each facet represents a predator, with color intensity indicating the number of records. Absence of a country from the map signifies no recorded predation for that predator.

### Computing Predation Preference and Probability of Mammal Species

Predicted predation probability was computed using bootstrapping paired with optimized XGBOOST models. Each dataset was randomly split into training (60%) and test (40%) sets, 250 times leading for each dataset to 250 different test-training combinations. XGBOOST models were computed using hyperparameter optimization (step size, max depth, max iterations) to minimize Logloss for classification (prey category) on the test set. The prediction was based for each species i on its traits using the following formula:

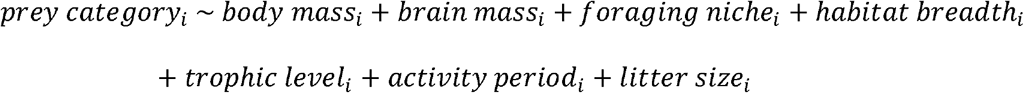

The optimization was done by grid searching on the following range of hyperparameters:

1. The step size of each boosting step (= 5, 6, 7 or 8).
2. The maximum depth of the tree (= 0.1 or 0.01).
3. The maximum number of iterations (= 10, 25, 50, 100 or 200).

Post-optimization probability for species not included in the training sets were calculated using the optimized model. Averaging probabilities per prey across the 250 test-training combinations yielded the final predictions per species for each dataset.

The importance of each trait in classifying prey species was assessed for each of the 9 predators based on the mean absolute SHAP value (mean(|SHAP value|)). For each predator, the average of the mean(|SHAP value|) was computed based on the 250 combinations generated for the bootstrapping. The calculation was performed using the sv_importance function from the shapviz R package (Mayer, 2024), which facilitates the interpretation of model predictions by quantifying the contribution of each feature. This approach considered all observations within each dataset because of the small sample sizes involved. To identify patterns of trait importance across the different predators, a principal component analysis (PCA) was conducted on the mean(|SHAP value|) of traits that exhibited variability (i.e., not exclusively null values across all predators). The PCA was executed using the prcomp function. The selection of the number of components to retain from the PCA was informed by the elbow method, which involves plotting the percentage of total variance explained by each component against the number of components. This method aids in identifying the point at which the addition of more components ceases to provide substantial additional explanatory value, thereby determining the optimal number of principal components to include in further analyses.

### Measuring Model Quality

To assess the quality of binary classifiers, I first used the Matthews correlation coefficient (MCC) based on the comparison between observed and predicted prey categories of species in each dataset because of its largely better insightful information compared to other standard metrics as area under the ROC curve (Chicco & Jurman 2020, 2023). The Matthews correlation coefficient was computed as follows:

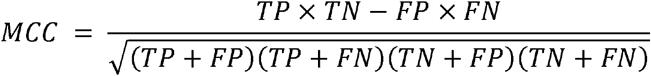

Where TP is the number of true positives, TN is the number of true negatives, FP is the number of false positives, and FN is the number of false negatives. For each of the 250 test-training combinations, one MCC was computed on the test set by comparing predicted and observed prey categories of this 30% remaining species of the original dataset. Consequently, the final value of MCC provided for each predator was the average MCC value across the 250 test-training combinations associated with the standard deviation.

The correlation between predicted probabilities and observed prey preference were assessed using the Spearman’s rank correlation coefficient between the predicted probability value and the observed predator preference with the cor function.

### Null Models

To ascertain the predictive power of traits for predation predictions, null models were created by shuffling predation preferences within the predator datasets with the sample R function, generating 99 null datasets per predator. The same quality metrics as for the actual datasets were applied. The efficacy of the models was evaluated by comparing the proportion of models with superior metrics against the null dataset outcomes.

## Results

### Binary classifier performances

The binary classifiers exhibited relatively good predictive capabilities across the nine predator species according to the Matthew correlation coefficient (MCC) values. The bootstrapping provided an average MCC of 0.35, with a median value of 0.30. This ranged from a minimal value of 0.22 for *Canis aureus* to a maximal value of 0.49 for *Lynx lynx*, indicating significant variability in predictive performance across the species (see Fig. 2, Table 1). Predators were clustered into two distinct groups based on their average MCC values: the first group, comprising *Canis aureus, Martes martes, Martes foina, Canis lupus*, and *Vulpes vulpes*, had MCC values ranging from 0.22 to 0.30; the second group, including *Felis silvestris, Genetta genetta, Neovison vison*, and *Lynx lynx*, exhibited higher MCC values between 0.44 and 0.49. The standard deviation of MCC values among all predators varied between 0.14 and 0.23.

**Table 1.** summarizes the model characteristics for each predator, including functional richness of the diet, calculated using the Hill number (Chiu & Chao, 2014) via the hillR package from traits for predation prediction, and adult body mass in grams, derived from a combined database (Soria et al., 2021).

| Scientific name | Exact correlation | Average value of MCC | Standard deviation of MCC | Number of predation record with preference | Functional richness of the diet | Body mass of adult (g) |
| --- | --- | --- | --- | --- | --- | --- |
| *Canis aureus* | 0.43 | 0.22 | 0.21 | 168 | 37.11 | 11000.00 |
| *Canis lupus* | 0.48 | 0.29 | 0.15 | 806 | 125.62 | 29190.76 |
| *Felis silvestris* | 0.56 | 0.44 | 0.17 | 172 | 204.33 | 5036.54 |
| *Genetta genetta* | 0.63 | 0.45 | 0.23 | 120 | 40.77 | 1866.67 |
| *Lynx lynx* | 0.60 | 0.49 | 0.15 | 315 | 131.11 | 21150.00 |
| *Martes foina* | 0.45 | 0.28 | 0.15 | 271 | 242.19 | 1700.00 |
| *Martes martes* | 0.39 | 0.25 | 0.17 | 207 | 115.08 | 1299.99 |
| *Neovison vison* | 0.52 | 0.45 | 0.19 | 103 | 45.17 | 899.75 |
| *Vulpes vulpes* | 0.46 | 0.30 | 0.14 | 800 | 436.15 | 4580.30 |

**Figure 2.**
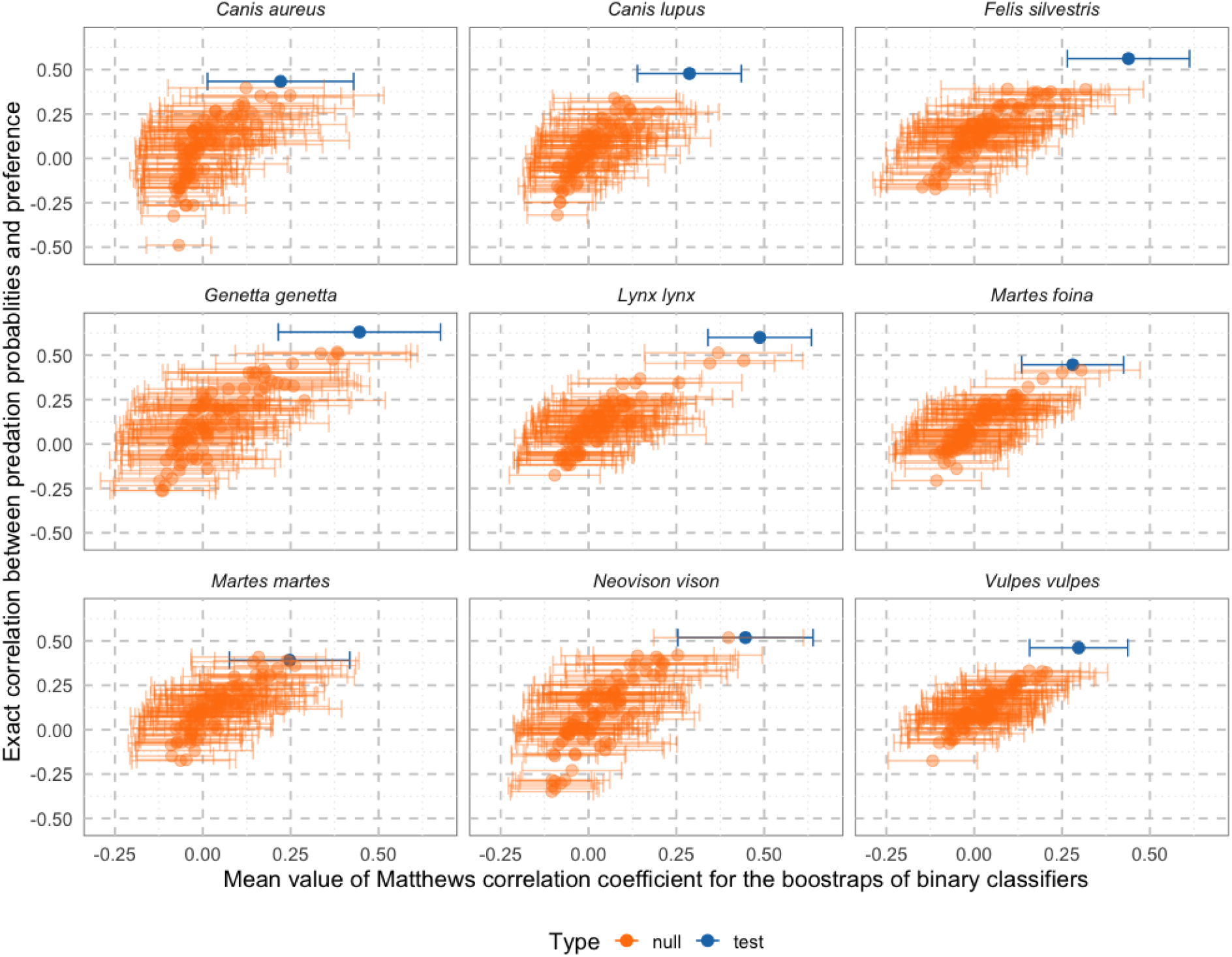
illustrates the correlation between predicted probabilistic predation and observed predation preferences for nine predators, factoring in prediction quality. Correlation was assessed using Spearman’s correlation coefficient, while model quality was gauged with the Matthews Correlation Coefficient (MCC). For MCC, points denote the average from 250 bootstrapped test-training dataset combinations, with brackets indicating variability. Blue points and brackets depict the test model, based on data from the CARNIDIET database, and orange points and brackets represent 99 models with CARNIDIET data randomly shuffled for each predator dataset.

Comparative analysis with null models highlighted that, with the exception of *Canis aureus, Martes martes*, and *Martes foina*, all species’ average MCC values were consistently higher when assessed with the original data rather than with their null counterparts. For the aforementioned three species, only a single null model outperformed the original data in terms of MCC value (Fig. 2).

Across all nine predators, the analysis identified common traits that accurately distinguished between prey and non-prey. However, the significance of these traits, as quantified by the mean absolute SHAP value, varied across the species (Fig. 3). A principal component analysis (PCA) of the mean absolute SHAP values for all traits used in the prediction models, capturing 48% of the total variance within the first two dimensions, revealed a single cluster. This cluster included *Canis aureus, Martes martes*, and *Vulpes vulpes*, indicating that for these species, traits similarly influenced the differentiation between predicted prey and non-prey. In contrast, the remaining species exhibited distinct patterns in their mean absolute SHAP values (Fig. 4).

**Figure 3.**
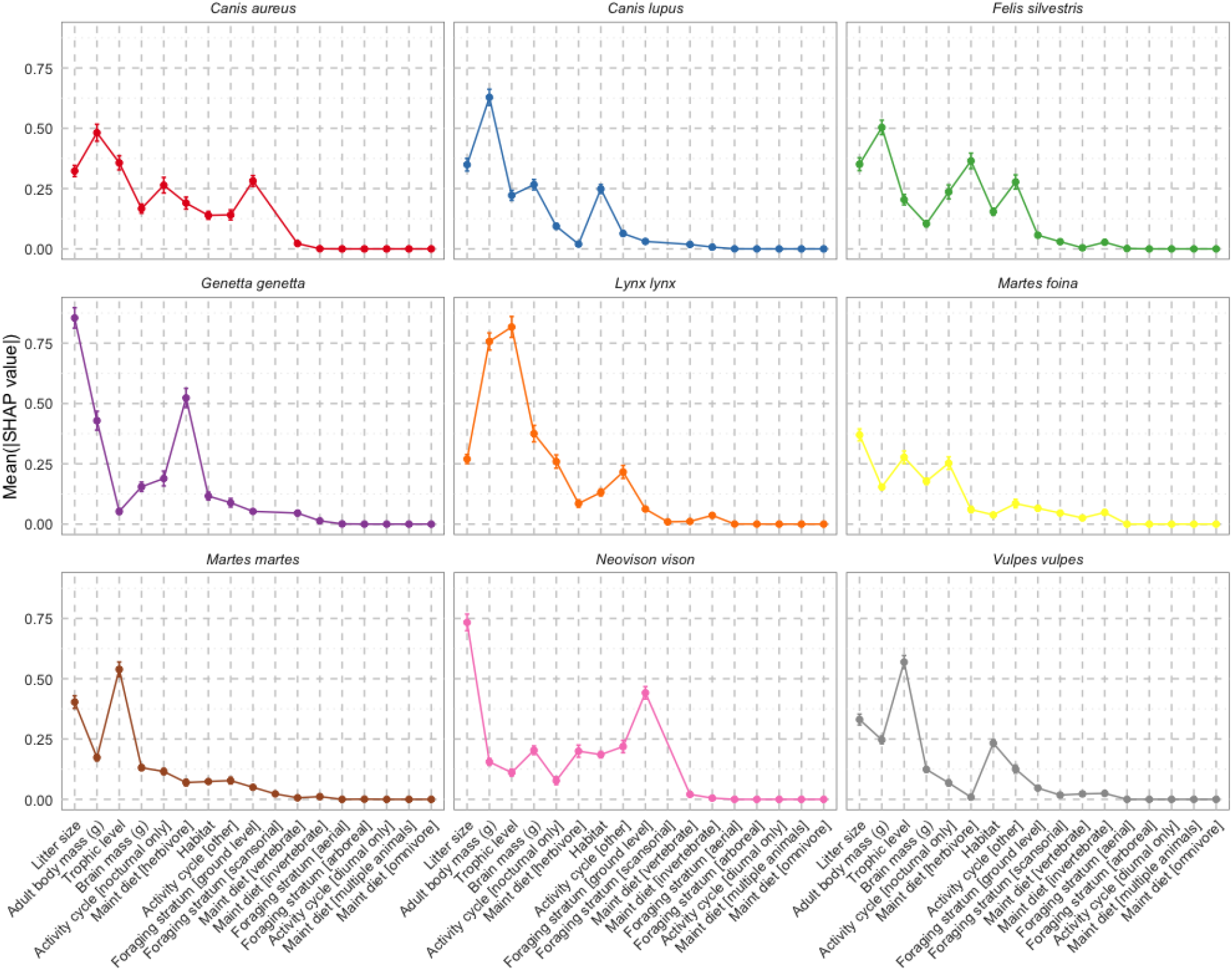
displays the significance of various traits in accurately predicting prey species for nine predators, based on the mean absolute SHAP value. Points indicate the average across 250 bootstrapped test-training dataset combinations, with brackets showing the standard deviation. A higher value signifies greater importance of the trait in determining whether a species is prey.

**Figure 4.**
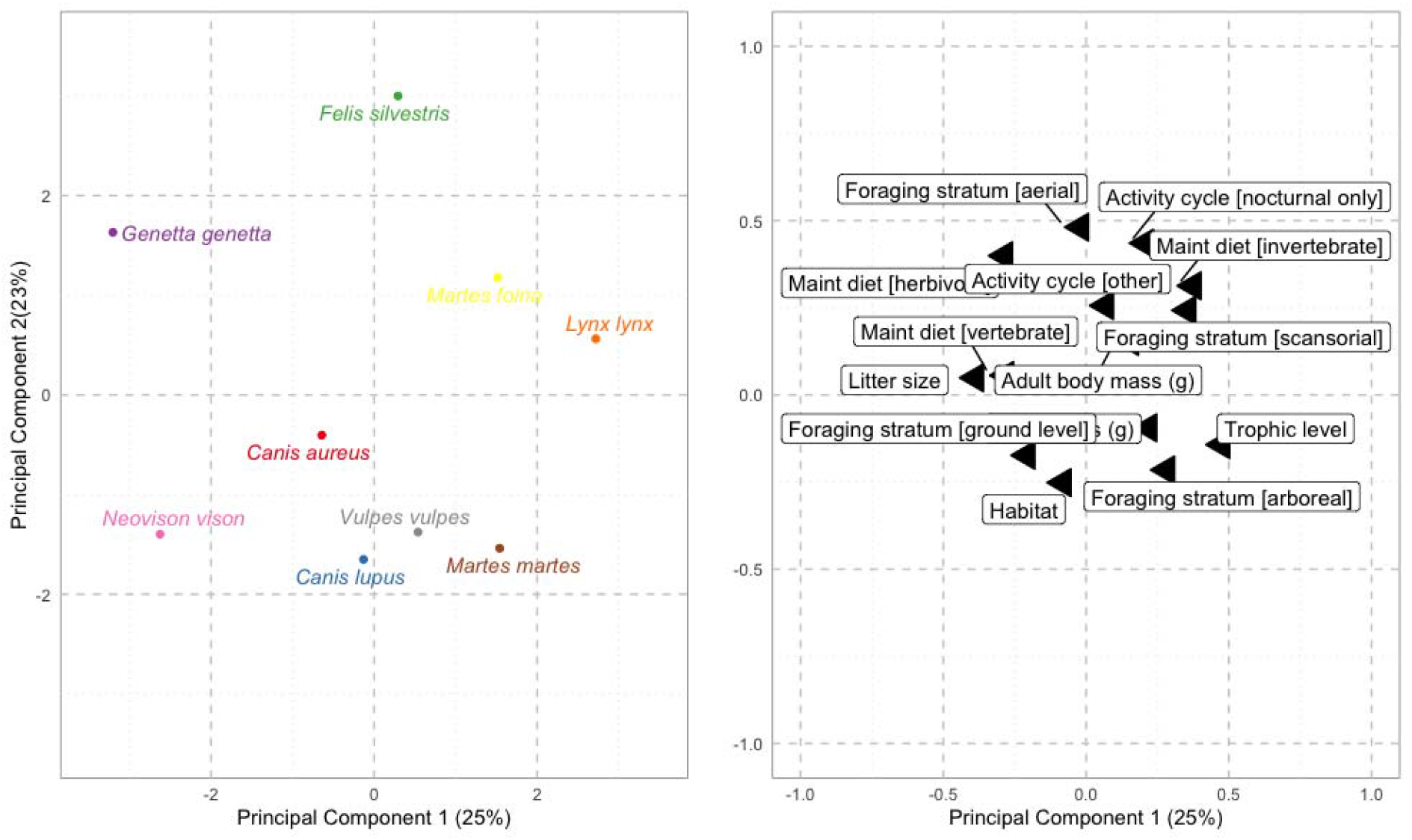
illustrates the differential impact of traits on predicting prey species, analyzed through principal component analysis (PCA) based on mean absolute SHAP values for traits showing variability across nine predators. Two components were chosen using the elbow method. The left panel displays each predator within the first two principal components, explaining 48% of the total variance, while the right panel correlates traits with their positions in this two-dimensional space.

### Predicted probabilities vs observed preferences

For the nine predators, the correlation between predicted predation probabilities and observed prey preferences showed a mean of 0.50 and a median of 0.48, with the range extending from 0.39 (*Martes martes*) to 0.63 (*Genetta genetta*) (Fig. 1, Table 1). Except for *Martes martes*, which had one null model showing a higher correlation, predictive models consistently outperformed null models (Fig. 1).

A detailed analysis revealed that lower predicted probabilities matched lower observed values closely, but higher probabilities exhibited more variability (Fig. 5). Notably, some species with high predicted probabilities were not prey according to the original dataset. Additionally, the observed probability range for predators like *Felis silvestris, Martes foina*, and *Vulpes vulpes* did not include any species with a predation probability below 0.2, unlike other predators, which had at least one species below a 0.1 probability.

**Figure 5.**
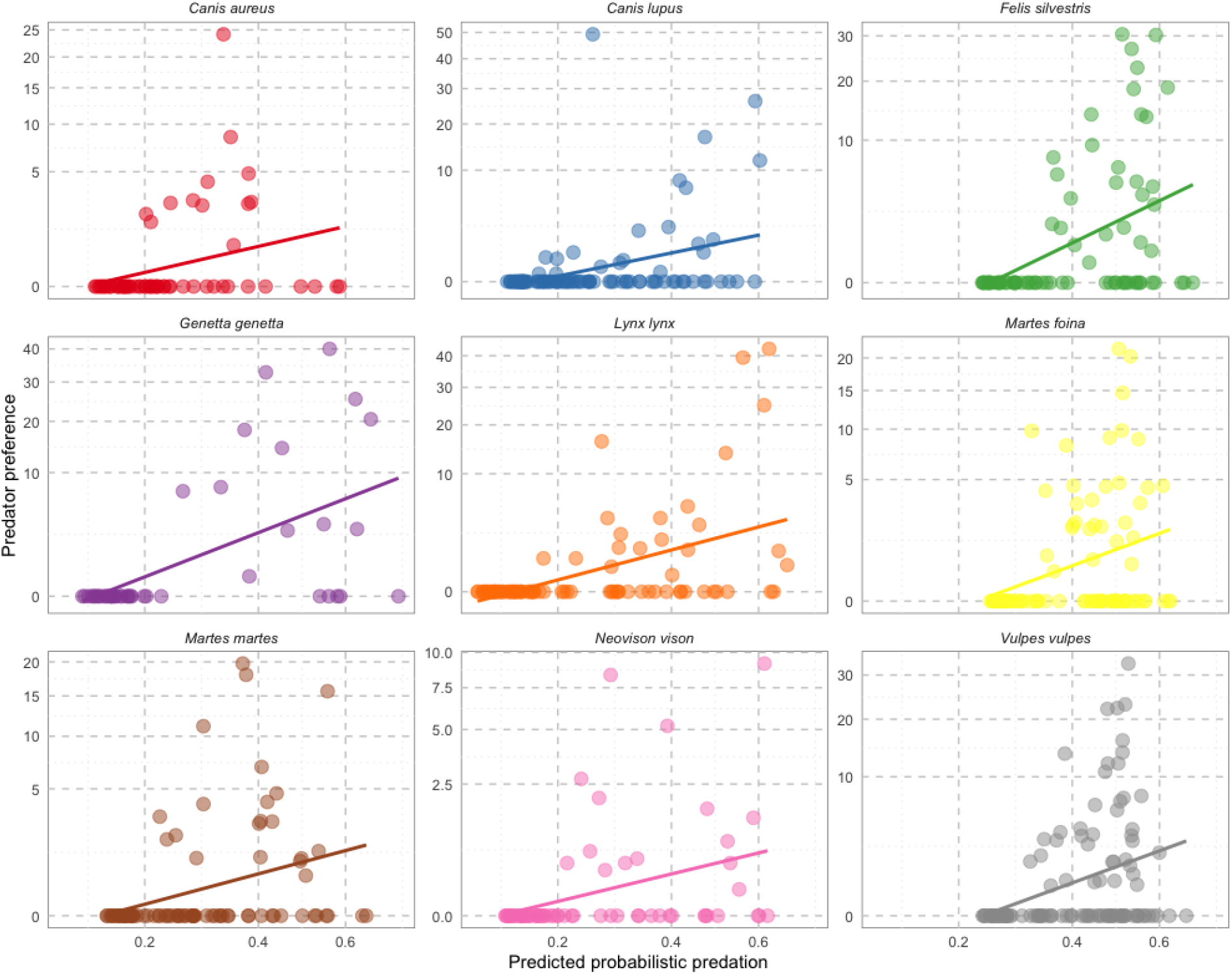
shows the linear correlation between predicted probabilistic predation and observed predation preference. Each point represents a mammal species whose geographical range intersects with at least one recorded predator location in the CARNIDIET database. The line depicts the linear regression between predicted probabilistic predation and observed predation preference.

## Discussion

This study demonstrates that the predicted probability of predation derived from binary classifiers can offer valuable insights into the intensity of prey-predator relationships. Utilizing the XGBOOST algorithm, the binary classifier effectively predicted predation events among nine mammalian predators, as demonstrated by the Matthew correlation coefficient. This research not only confirms the classifier’s predictive capability but also highlights the significant correlation between predicted probabilistic predation and observed prey preferences. These findings suggest that probabilistic predation models could enhance the accuracy of trophic network analyses by providing a more nuanced understanding of these relationships.

Further, the study highlights the importance of selecting informative species traits, as indicated by the stark contrast in performance between our models and null models across all predators. This underscores the predictive value of well-chosen traits in understanding prey-predator dynamics.

The efficacy of binary classifiers in predicting predation, utilizing both regression and tree-based algorithms, is now widely recognized across diverse trophic levels and taxa, incorporating traits and phylogeny (Caron et al., 2022; Desjardins-Proulx et al., 2017; Llewelyn et al., 2022, 2023). Despite a small sample size, this study’s classifiers maintained robust performance, as validated by the Matthew correlation coefficient and further supported through bootstrapping variability analysis.

Analysis of SHAP values revealed that, despite differing magnitudes, similar traits were consistently associated with predation across the nine studied predators, all belonging to the Carnivora order. However, individual species demonstrated unique patterns in how these traits influenced predation predictions, with notable exceptions including *Canis aureus, Martes martes*, and *Vulpes vulpes*. Given the limited scope of predator species analyzed, broader conclusions regarding model influences remain speculative, suggesting a need for future research into how predator characteristics and data quality might affect classifier performance.

Moreover, the correlation between binary classifier-derived predation probabilities and observed predation preferences was substantively strong across all species, with Spearman’s correlation coefficients ranging from 0.39 to 0.63. This finding underscores the potential for probabilistic models to accurately reflect predation preferences, both ranging between 0 and a maximum value, even with relatively small datasets and broad-scale predictions (even if that we know that trophic relationships are not stable across space and time, see for example Castañeda et al. (2022) for foxes and Jensen et al. (2022) for coyotes). Strikingly, I observed that for all predators there were numerous species with a high probability of predation but not observed initially as prey. This apparent paradox can be explained by two hypotheses, although it’s not possible to discern between the two. On one hand, the observed data may have an observation bias, and the predation probability would ‘correct’ this bias, because according to the species’ traits, there is little chance that it is not prey. On the other hand, the model may be based on some traits that do not capture all the dimensions that make a species prey, and thus biases its predation probability for certain species. So, the more exhaustive the binary interaction database, the weaker the first hypothesis becomes, and vice versa. Reducing to zero the probability of predation for species for which no predation record exists could also be considered, but this action would have to be based on solid hypotheses regarding the model’s misassignment, such as a very low sampling bias.

This preliminary study, focusing on a selected group of predators and mammalian prey, lays the groundwork for extending probabilistic predation models to encompass comprehensive trophic networks. Although binary interaction data are increasingly available, corroborating these with prey preference data remains a challenge. Future research should prioritize the compilation of prey preference data, facilitated by advances in biodiversity sampling technologies such as camera trapping and eDNA analysis (Strydom et al., 2021).

As trophic interactions gain prominence in biogeography and macroecology, understanding and modeling these relationships become crucial for elucidating fundamental patterns and addressing conservation challenges (O’Connor et al., 2020; Strona & Bradshaw, 2022; Thuiller et al., 2023). Transitioning from binary to probabilistic interaction models represents a significant opportunity to refine our analyses and explore novel outcomes in ecological research.

## Data availability

All data and codes are available in github:

https://github.com/MartPhilip/Advancing_ecological_networks

